# The role of the cingulate cortex in the generation of motor tics and the experience of the premonitory urge-to-tic in Tourette syndrome

**DOI:** 10.1101/2020.01.29.924399

**Authors:** Stephen R. Jackson, Hilmar P. Sigurdsson, Katherine Dyke, Maria Condon, Georgina M. Jackson

## Abstract

Tourette syndrome (TS) is a neurological disorder of childhood onset that is characterised by the occurrence of motor and vocal tics. TS is associated with cortical-striatal-thalamic-cortical circuit [CSTC] dysfunction and hyper-excitability of cortical limbic and motor regions that are thought to lead to the occurrence of tics. Individuals with TS often report that their tics are preceded by ‘premonitory sensory/urge phenomena’ (PU) that are described as uncomfortable bodily sensations that precede the execution of a tic and are experienced as a strong urge for motor discharge. While the precise role played by PU in the occurrence of tics is largely unknown, they are nonetheless of considerable theoretical and clinical importance as they form a core component of many behavioural therapies used in the treatment of tic disorders. Recent evidence indicates that the cingulate cortex may play an important role in the generation of PU in TS, and in ‘urges-for-action’ more generally. In the current study we utilised voxel-based morphometry (VBM) techniques, together with ‘seed-to-voxel’ structural covariance network (SCN) mapping, to investigate the putative role played by the cingulate cortex in the generation of motor tics and the experience of PU in a relatively large group of young people with TS. Whole-brain VBM analysis revealed that TS was associated with clusters of significantly reduced grey matter volumes bilaterally within: the orbito-frontal cortex; the cerebellum; and the anterior and mid cingulate cortex. Similarly, analysis of SCNs associated with bilateral mid- and anterior-cingulate ‘seed’ regions demonstrated that TS is associated with *increased* structural covariance primarily with the bilateral motor cerebellum; the inferior frontal cortex; and the posterior cingulate cortex.

## Introduction

Tourette syndrome (TS) is a neurological disorder of childhood onset that is characterised by the presence of chronic vocal and motor tics (Cohen, Leckman, & Bloch, 2013). Tics are involuntary, repetitive, stereotyped behaviours that occur with a limited duration (Cohen et al., 2013). Motor tics can be simple or complex in appearance, ranging from repetitive movements to coordinated action sequences. Verbal tics can consist of repetitive sounds, words or utterances, the production of inappropriate or obscene utterances, or the repetition of another’s words. Tics occur in bouts, typically many times in a single day, and are the most common form of movement disorder in children.

Individuals with TS perceive a relatively constant demand to suppress their tics, particularly in social situations, and while the voluntary suppression of tics is possible in many cases, they typically report that it can be uncomfortable and stressful to suppress tics, and that the urge to tic becomes uncontrollable after a period of suppression. Importantly, the majority of individuals with TS report that their tics are often preceded by ‘premonitory sensory/urge phenomena’ (PU) that are described as uncomfortable cognitive or bodily sensations that occur prior to the execution of a tic and are experienced as a strong urge for motor discharge (Cohen et al., 2013. Individuals who experience PU often report that: these experiences are more bothersome than their tics; that expressing their tics give them relief from, and temporarily *abolishes*, their PU; and that they would not exhibit tics if they did not experience PU (Cavanna, Black, Hallett, Voon, 2017). For this reason, it has been proposed that PU should be considered as the driving force behind the occurrence of tics, and that tics are a learnt response to the experience of PU (Cavanna et al., 2017). PU are of particular clinical importance as they form a core component of behavioural therapies that are currently used in the treatment of tic disorders (Cohen et al., 2013).

Our understanding of PU and their relationship to tics is currently limited, and there are grounds for thinking that the occurrence of tics and the occurrence of PU are independent processes or only loosely associated. First, not all individuals with TS report experiencing PU. In particular, children under 10 years of age, who present with simple tics, do not typically report being aware of PU (Cohen et al., 2013). Second, tics have been observed during sleep, including slow-wave sleep, indicating that at least some tics are involuntary (Cohrs et al., 2001). Third, the occurrence of tics - and an individual’s ability to suppress them - may occur independently of the awareness of PU (Ganos Kahl, Schunke, Kuhn, Haggard, Gerloff, el al., 2012). Finally, the generation of tics and the genesis of PU in TS have been linked to different brain networks (Bronfeld, Israelashvili, & Bar-Gad, 2013; Conceicao, Dias, Farinha, & Maia, 2017; Jackson, Parkinson, Kim, Schuermann, & Eickhoff, 2011; McCairn, Iriki, & Isoda, 2013). Previous studies have indicated that the urge-for-action more generally may activate a common set of brain areas across a wide range of behavioural domains (e.g., the urge to blink, the urge to yawn, the urge to micturate, the urge to scratch an itch, etc.), that includes the urge to tic in Tourette syndrome (TS) (Jackson, Parkinson, Kim, et al., 2011). Jackson and colleagues conducted a quantitative meta-analysis of functional brain imaging studies that investigated the ‘urge-for-action’ associated with everyday behaviours such as yawning, swallowing, and micturition, and demonstrated that the right anterior insula and the mid-cingulate cortex were the only regions consistently activated across brain imaging studies associated with the perception of the urge for action in different behavioural domains (Jackson, Parkinson, Kim, et al., 2011). Importantly, these authors proposed that the right insula and mid-cingulate cortex play a central role in a neural circuit that represents bodily sensations, generates urges for action, selects an action based upon an estimation of the likely outcomes associated with that action, and determines whether the conditions giving rise to the urge-for-action have been resolved once an action has been initiated.

Consistent with this proposal, functional brain imaging studies indicate that brain activity within the mid-cingulate cortex increases one second prior to tic execution in individuals with TS (Bohlhalter et al., 2006; Neuner et al., 2014) and structural brain imaging studies demonstrate that there are alterations in grey matter (GM) volume throughout mid and anterior cingulate cortex that are correlated with clinical measures of tic severity (for a recent review see O’Neill, Piacentini & Peterson, 2019). This has led some authors to conclude that the mid cingulate cortex plays an important role in the generation/representation of premonitory urges (e.g., O’Neill et al., 2019) whereas others have speculated that the role of the cingulate motor area may be to select/generate a particular action in response to PU that may be primarily generated elsewhere, most likely within the anterior insula (Jackson, Parkinson, Kim, et al., 2011). This latter view is consistent with recent studies demonstrating that while electrical stimulation of medial wall regions of cortex, including the mid cingulate cortex, was sufficient to induce movements, including goal-directed actions, there was no evidence that electrical stimulation of this region induced a phenomenological experience of an ‘urge-to-move’ (Caruana, Gerbella, Avanzini, Gozzo, Pelliccia, Mai, Abdollahi, Cardinale, Sartori, Lo Russo, Rizzolatti, 2018; Trevisi, Eickhoff, Chowdhury, Jha, Rodionov, Nowell, Miserocchi, McEvoy, Nachev, Diehl, 2018; although see Fried, Katz, McCarthy, Sass, Williamson, Spencer, Spencer, 1991, for an alternative report that electrical stimulation of the posterior SMA can induce the experience of an ‘urge-to-move’). In the current study we focus specifically on the relationship between the cingulate cortex, measured using structural magnetic resonance imaging together with the analysis of structural covariance networks, and clinical measures of tic severity and PU.

Human brain imaging studies have identified a number of functional brain networks, often referred to as ICNs (intrinsic cortical networks) that reflect correlated brain activity across anatomically separate brain areas. Recent evidence indicates that these networks are dominated by common organisational principles and stable features, and may largely reflect enduring individual characteristics, including the consequence of brain health conditions (Gratton et al., 2018). Similarly, neuroimaging studies have repeatedly demonstrated covariance of cortical thickness or grey matter (GM) volume over widespread, distributed, brain regions; and these structural covariance networks (SCNs) have also been shown to be highly heritable and to reflect differences in age and disease status (Alexander-Bloch, Giedd, & Bullmore, 2013).

It has been proposed that structural covariance between brain regions may likely reflect brain areas that are functionally co-active and exhibit common patterns of maturational change - including shared long-term trophic influences; shared patterns of gene co-expression (Romero-Garcia, Whitaker, Vasa, Seidlitz, Shinn, Fonagy, et al., 2018; Zielinski, Gennatas, Zhou, & Seeley, 2010), and are selectively vulnerable to specific brain health conditions (Seeley, Crawford, Zhou, Miller, & Greicius, 2009). Importantly, recent studies have demonstrated that SCNs closely mirror the functional ICNs revealed using resting-state functional magnetic resonance imaging [fMRI] (Kelly, Toro, Di Martino, Cox, Bellec, Castellanos, et al., 2012; Seeley et al., 2009) and co-degenerate in distinct human neurodegenerative conditions (Cauda, Nani, Manuello, Premi, Palerno, Tatu, et al., 2018; Seeley et al., 2009). This suggests that analysis of SCNs, while currently under-utilised to study brain networks in neurodevelopmental conditions, may be a particularly useful method for investigating alterations in brain network development in children and adolescents for whom the use of conventional fMRI approaches is especially challenging. In this study we chose to investigate specifically how SCNs associated with different functional regions of the bilateral cingulate cortex may be altered in children and adolescents with Tourette syndrome (TS) relative to a group of typically developing individuals.

## Materials and Methods

This study was approved by the University of Nottingham, School of Psychology, ethical review committee. Written informed consent was acquired from all participants and where appropriate from their parents/caregivers. No part of the study procedures or analyses were pre-registered prior to the research being conducted. We report how we determined our sample size, all data exclusions (if any), all inclusion/exclusion criteria, whether inclusion/exclusion criteria were established prior to data analysis, all manipulations, and all measures in the methods below. Finally, the conditions of our ethics approval do not permit public archiving of individual MRI data or clinical biographical data.

### Participants

In total 76 volunteers took part in this study: 39 had a confirmed diagnosis of TS (TS group) and 37 formed our control group (CS group) of age- and sex-matched, typically developing, individuals with no history of neurological disorders. The TS group were recruited either from the Child and Adolescent Psychiatry Clinic at the Queens Medical Centre in Nottingham or by advertising through the Tourettes Action charity or regional TS support groups. The CS group were recruited from local schools, by local advertising, and recruitment at science fairs. All volunteers were provided with a small inconvenience allowance for their participation.

After magnetic resonance imaging (MRI) the scans from twelve participants were found to be un-usable and data from these individuals were excluded from further analyses. The participants who remained included 28 individuals in the TS group (3 females; mean age 14.62 ± 3.4 years) and 36 controls (3 females; mean age 14.38 ± 3.2 years). 10 individuals with TS had a confirmed or suspected clinical diagnosis of a co-occurring neuropsychiatric condition in addition to their TS (attention deficit/hyperactivity disorder [ADHD] = 2; obsessive-compulsive disorder [OCD] =2; and autism spectrum disorder [ASD] = 6). 10 patients were medicated at the time of scanning. Details of the TS group are reported in Table 1.

**Table 1.** Details of TS participants

| ID | Gender | Age | IQ | TIV | YGSS | Motor | Phonic | Impairment | PUTS | Comorbidity |
| --- | --- | --- | --- | --- | --- | --- | --- | --- | --- | --- |
| TS013 | M | 19.29 | 135 | 1672.99 | 32/100 | 18/25 | 9/25 | 5/50 | 17/36 | - |
| TS030 | M | 17.03 | 103 | 1534.19 | 23/100 | 12/25 | 11/25 | 0/50 | 10/36 | - |
| TS031 | M | 20.12 | 111 | 1393.68 | 67/100 | 21/25 | 16/25 | 30/50 | 22/36 | - |
| TS034 | M | 16.56 | 118 | 1572.93 | 0/100 | 0/25 | 0/25 | 0/50 | 34/36 | - |
| TS043 | M | 14.09 | 123 | 1521.32 | 63/100 | 23/25 | 20/25 | 20/50 | 21/36 | - |
| TS048 | M | 16.82 | 118 | 1560.25 | 19/100 | 9/25 | 0/25 | 10/50 | 16/36 | - |
| TS058 | M | 10.04 | 96 | 1488.58 | 10/100 | 6/25 | 4/25 | 0/50 | 16/36 | ADHD |
| TS061 | M | 11 | 99 | 1604.86 | 26/100 | 12/25 | 9/25 | 5/50 | 23/36 | - |
| TS066 | M | 16.16 | 102 | 1579.58 | 42/100 | 16/25 | 16/25 | 10/50 | 19/36 | OCD |
| TS069 | M | 13.47 | 133 | 1678.11 | 19/100 | 8/25 | 6/25 | 5/50 | 9/36 | - |
| TS071 | M | 15.61 | 118 | 1545.65 | 31/100 | 10/25 | 11/25 | 10/50 | 21/36 | - |
| TS075 | M | 16.25 | 112 | 1602.50 | 30/100 | 12/25 | 8/25 | 10/50 | 17/36 | - |
| TS084 | F | 22.78 | 126 | 1495.58 | 25/100 | 8/25 | 7/25 | 10/50 | 33/36 | ASD, GAD,<br>MDD |
| TS087 | M | 12.79 | 119 | 1509.92 | 30/100 | 15/25 | 10/25 | 5/50 | 23/36 | - |
| TS088 | M | 13 | 109 | 1527.10 | 71/100 | 20/25 | 21/25 | 30/50 | 25/36 | - |
| TS094 | F | 18.75 | 116 | 1292.99 | 32/100 | 12/25 | 15/25 | 5/50 | 21/36 | ASD |
| TS095 | M | 14.49 | 115 | 1452.84 | 23/100 | 13/25 | 0/25 | 10/50 | 30/36 | - |
| TS096 | M | 8.61 | 126 | 1544.50 | 23/100 | 10/25 | 3/25 | 10/50 | 18/36 | ADHD |
| TS097 | M | 15.28 | 112 | 1494.73 | 33/100 | 10/25 | 13/25 | 10/50 | 30/36 | OCD |
| TS101 | M | 10.48 | 129 | 1699.19 | 40/100 | 16/25 | 14/25 | 10/50 | 20/36 | ASD |
| TS102 | M | 15 | 111 | 1552.28 | 13/100 | 8/25 | 0/25 | 5/50 | 17/36 | - |
| TS103 | M | 18.96 | 113 | 1510.52 | 64/100 | 22/25 | 22/25 | 22/50 | 17/36 | - |
| TS105 | M | 11.35 | 115 | 1596.10 | 42/100 | 13/25 | 15/25 | 20/50 | 19/36 | - |
| TS110 | M | 13.22 | 85 | 1631.39 | 9/100 | 9/25 | 0/25 | 0/50 | 10/36 | ASD |
| TS113 | M | 13.1 | 92 | 1580.11 | 67/100 | 19/25 | 18/25 | 30/50 | 20/36 | ASD |
| TS119 | M | 9.67 | 107 | 1470.05 | 29/100 | 13/25 | 11/25 | 5/50 | 14/36 | - |
| TS127 | F | 13.01 | 75 | 1355.28 | 27/100 | 10/25 | 7/25 | 10/50 | 12/36 | ASD |
| TS139 | M | 12.36 | 100 | 1710.96 | 60/100 | 15/25 | 15/25 | 30/50 | 32/36 | - |

**Abbreviations.** Motor, phonic and impairment scores were measured using the YGTSS (Yale Global Tic Severity Scale); PUTS (Premonitory Urge for Tics Scale). TIV (Total intracranial volume). ADHD (attention deficit/hyperactivity disorder); OCD (obsessive compulsive disorder); ASD (autism spectrum disorder); MDD (major depressive disorder); GAD (generalized anxiety disorder). YGTSS scores were measured on the day of testing.

### Diagnosis, symptom severity and screening

Diagnosis of TS was confirmed by an experienced clinician. In addition, all participants underwent comprehensive screening for current symptoms of TS by a highly experienced and trained research nurse/researcher. Measures of the current severity of tics were obtained using the Yale Global Tic Severity Scale (YGTSS) (Leckman et al., 1989). The YGTSS is a semi-structured clinician-rated measure assessing the nature of motor and vocal tics present over the past week. The YGTSS is a commonly used clinical assessment scale and has been found to have good psychometric properties (Leckman et al., 1989). It consists of three subscales: impairment rating, motor tic rating and vocal tic rating. Motor and vocal tic ratings are made up of the composite answers from questions relating to number, frequency, intensity, complexity and interference of tics reported in the previous week and observed during the interview. The current frequency and severity of premonitory sensory/urge phenomena [PU] was measured using the Premonitory Urge for Tics Scale (PUTS) (Woods, Piacentini, Himle, & Chang, 2005). The PUTS is a self-report measurement where items assess the intensity and frequency of PSP (on a scale of 1 – 4). 9 of the 10 items on the PUTS scaled were scored based on recommendation, and thus scores could range from 9 to 36 (Woods et al., 2005). Participants were screened for any indication of symptoms of ADHD, OCD and Autism using the Connors-3 Parent Report (Conners, 2008), Children’s Yale-Brown Obsessive-Compulsive Scale (CY-BOCS) (Scahill et al., 1997) and Social Communication Questionnaire (SCQ) (Berument, Rutter, Lord, Pickles, & Bailey, 1999), respectively. Based on these measures, a further eight patients were categorised as being at high risk of having OCD and/or ADHD. All participants also completed the Wechsler’s Abbreviated Scale of Intelligence (WASI-II) (Wechsler, 1999) used to assess intellectual ability. Two subtests were used (the verbal and performance subtests). Participants would be excluded from the study if their WASI score was < 70 (none were).

### Image acquisition

Whole-brain, high-resolution, T1-weighted structural MRI brain images were acquired for each participant. Scanning was conducted at the Sir Peter Mansfield Imaging Centre (SPMIC), Nottingham, UK using a 3T Philips Achieva MRI scanner with a 32-channel SENSE head-coil and running a MPRAGE sequence (180 contiguous axial slices, 8.6 ms repetition time [TR], 4.0 ms echo time [TE], 256 × 224 × 180 matrix size, 1×1×1 mm raw voxel size, and a scan duration of 225 seconds). Prior to acquisition, participants were asked to lie as still as possible with their eyes open. Foam padding was added for extra stability and to reduce head movements. All participants wore also noise-cancelling headphones.

### MR data pre-processing

Pre-processing of all MRI images was accomplished using SPM12 and the Computational Anatomy Toolbox (CAT12; http://www.neuro.uni-jena.de/cat/). First, raw structural T1-w MRI scans were oriented to have the origin lying on the AC-PC line using automated registration. Intensity normalisation, bias and noise-correction was conducted using the Spatially Adaptive Non-Local Means (SANLM) tool in CAT12 and the images were spatially normalised using DARTEL (affine and non-linear registration, [Ashburner, 2007]) to standard space and segmented into different tissue classes: grey matter (GM), white matter (WM) and cerebrospinal fluid (CSF). The images were then modulated - which involves scaling by the amount of contraction done during the normalisation step – to ensure that GM in the normalised images remains the same as in the raw images. Finally, all de-noised, normalised, segmented and modulated GM maps were smoothed using an 8-mm full-width at half maximum (FWHM) Gaussian kernel. Prior to pre-processing, all scans were visually inspected and images with any visible artifact were excluded. Total intracranial volume (TIV) was estimated from all subjects. The GM maps were co-registered to the AAL2 atlas to enable an anatomically defined region-of-interest (ROI) for the bilateral mid and anterior cingulate cortex to be generated.

### Structural covariance

Three separate functionally-defined bilateral ROIs were created based upon the control data published by Balsters and colleagues (Balsters, Mantini, Apps, Eickhoff, Wenderoth, 2016: connectivity based probability maps of the cingulate cortex for typically developing young adults are available for download at http://www.ncm.hest.ethz.ch/downloads/data.html). These functionally-defined ROIs consisted of: a bilateral posterior mid-cingulate region; a bilateral anterior mid-cingulate region; and a bilateral posterior anterior-cingulate region. Then, using the segmented whole-brain GM maps and a ‘seed-to-voxel' approach, we computed the structural covariance between the mean GM values for voxels within each of our empirically-defined cingulate (seed) ROIs and the GM values obtained for all voxels in the GM maps. This analysis yielded a covariance map for each group in which the value at each voxel reflected the cross-subject Pearson correlation coefficient between the mean GM value for the seed region (the respective cingulate cortex ROI) and the GM value at that particular voxel. The correlation coefficients were converted to Z scores using Fisher’s r-to-Z transformation and the whole-brain Z(r) maps for each group were then statistically compared at group-level using the following equation

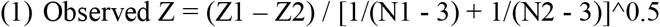

where, for each voxel: Z1 is the Z(r) value for that voxel for the TS group; Z2 is the Z(r) value for that voxel for the CS group; N1 the number of participants in the TS group; and, N2 the number of participants in the CS group. The computed Z-maps were corrected for multiple comparisons using FDR [p-FDR < 0.05] (Benjamini & Hochberg, 1995) and a cluster threshold of K_E_ ≥ 50 voxels was applied. Labelling of statistically significant clusters was accomplished using the Brain Anatomy Toolbox (Eickhoff et al., 2005). By definition, two regions ‘covary’ positively when increased GM values in one region is associated with increased GM values in another region. We defined *negative* covariance as an increase in GM values in one region that is associated with a reduction in GM values in a separate region.

## Results

### Preliminary analyses: group differences

The preliminary analyses of these data have been reported previously (Jackson, Loayza, Crighton, Sigurdsson, Dyke, Jackson (in press) but for completeness they are reported here. An independent samples t-test confirmed that there was no significant difference in age between the TS (mean =14.62 ± 3.43) and CS (mean = 14.38 ± 3.23) groups (t(62) = −0.28, p= 0.78). However, independent samples t-tests revealed that there was a significant between-group difference in total intracranial volume [TIV] (TS mean = 1544.97 ± 97.15; CS mean =1640.88 ± 158.71; t(62)=2.81, p=0.007), with controls having a higher TIV than individuals with TS, and a significant between-group difference in IQ (TS mean = 111.36 ± 13.93; CS mean = 118.58 ± 12.28; t(62)=2.20, p=0.03) with controls exhibiting a slightly higher average IQ. It should be noted however that both groups exhibited above-average IQ scores. For the whole-brain VBM and structural covariance analyses reported below the adjusted grey matter volumes were used after co-varying for age, sex, IQ and TIV.

### Exploring differences in total intracranial volume

Preliminary analysis of the anatomical MRI images revealed a significant between-group difference in TIV, with the CS group exhibiting a significantly larger mean TIV value that the TS group. However, interpretation of this finding is challenging without further analyses due to differences in Age, Sex, and IQ between the groups, and because the TIV measure includes different tissue types (i.e., GM, WM, and CSF). To further explore this finding, and as this paper is concerned with GM morphometry, we calculated two additional measures: first, the total number of GM voxels within each GM map for each participant; and second, the average GM value across all of the GM voxels within each GM map for each participant. Furthermore, two TS participants who were each aged over 30 years of age were excluded from this analysis.

For each of these measurements we conducted a separate stepwise multiple regression analysis, with the following variables as predictors: Chronological age; Sex; IQ; Group (CS vs. TS); and the Group x Age interaction. Note that in each case, the initial order of entry was fixed with Chronological age entered first into the model first.

#### Analysis of total number of GM voxels

A scatterplot illustrating how the number of GM voxels in each GM map decreases as a function of Chronological age for each group is illustrated in Figure 1A. Inspection of this figure illustrates that while there is a small and only marginally significant decrease in number of GM voxels with age for the CS group (R^2^ = 0.11, p = 0.054), the decrease with age for the TS group is much steeper and statistically significant (R^2^= 0.48, p = 0.0001). The results of the stepwise multiple regression analysis demonstrated that Chronological age (t = −3.79, p < 0.0003) and the Age x Group interaction (t = 2.64, p < 0.01) were each independent and statistically significant and predictors of total number of GM voxels. The final regression model accounted for more than 24% of the variance in total number of GM voxels (F = 10.87, adjusted-R^2^ = 0.24, p < 0.0001). The results of this analysis confirm that after differences in chronological age have been taken into account, there is a large and statistically significant effect of group in the form of an Age x Group interaction, which indicates that the number of GM voxels decreases more steeply with age in the TS group.

**Figure 1:**
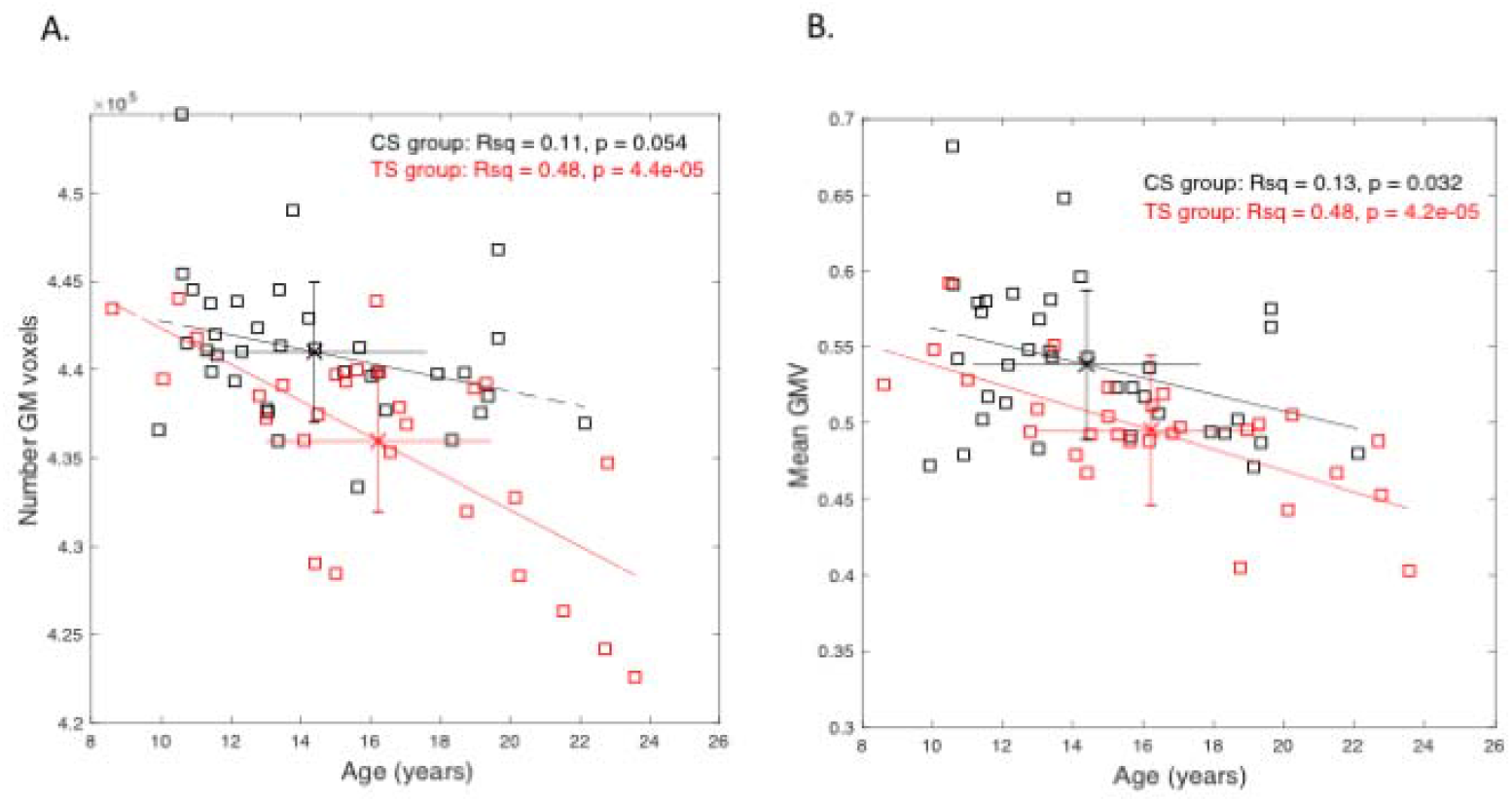
Illustrates associations between GM measures and chronological age for each group. **A**. Shows the association between the total number of GM voxels observed for each group as a function of chronological age. **B**. Shows the association between the mean GM values observed for each group as a function of chronological age. Error bars represent the standard deviation.

#### Analysis of average GM values

Figure 1B illustrates how the mean GM values within each GM map decreases as a function of Chronological age for each group. Inspection of this figure illustrates that while there is a decrease in mean GM values with age for both groups (CS group: R^2^ = 0.13, p = 0.05; TS group: R^2^ = 0.48, p = 0.0001), the mean GM value for the TS group appears substantially lower at each age than for the CS group. The results of the stepwise multiple regression analysis demonstrated that Chronological age (t = −4.05, p < 0.0001) and the Age x Group interaction (t = 3.48, p < 0.001) were each independent and statistically significant and predictors of the mean GM value in each GM map. The final regression model accounted for approximately 30% of the variance in mean GM value (F = 14.54, adjusted-R^2^ = 0.30, p < 0.0001).

The results of this analysis confirm that once differences in chronological age and sex have been taken into account, there remains a large and statistically significant effect of group, which indicates that the mean GM value is significantly reduced for the TS group at all ages examined.

Overall, these data indicate that GM differences in the TS group are unlikely to be due to between-group differences in chronological age, sex, or IQ, but instead are largely associated with the clinical diagnosis of TS.

### Whole-brain voxel-based morphometry analysis

The primary objective of this study is use voxel-based morphometry techniques together with ‘seed-to-voxel’ structural covariance network (SCN) mapping to investigate the role played by the mid and anterior cingulate cortex in the generation of motor tics and the experience of PU in a relatively large group of people with TS. However, for completeness, and given the preliminary analysis of whole-brain GM reported above, we now report the results of a whole-brain between-group VBM analysis.

The VBM analysis was carried out using the Computational Anatomy Toolbox (CAT12) and SPM12. An independent samples t-test was conducted to compare the GM maps of the CS and TS groups. Importantly, the following variables were modelled as covariates of no interest: chronological age; sex, TIV, and IQ. The resultant t-maps were then corrected using the CAT12 cluster correction function. The resulting clusters are presented in Table 2 below.

**Table 2:** Results of the whole-brain VBM analysis (independent groups t-test) for the CS > TS contrast.

| Cluster | Size | t-value | Region | Hemisphere | Peak MNI coordinates |  |  |
| --- | --- | --- | --- | --- | --- | --- | --- |
|  |  |  |  |  | X | Y | Z |
| 1 | 106 | 4.77 | middle frontal gyrus | L | -35 | 1 | 45 |
| 2 | 103 | 4.42 | anterior cingulate cortex | L | -5 | 38 | 14 |
| 3 | 35 | 4.20 | precentral gyrus | R | 37 | 1 | 47 |
| 4 | 71 | 3.85 | medial orbital gyrus | R | 15 | 54 | -14 |
| 5 | 17 | 3.75 | Hippocampus | R | 27 | -20 | -23 |
| 6 | 40 | 3.62 | supplementary motor cortex | L | -11 | 28 | 32 |

#### CS group > TS group contrast

The analysis revealed six clusters that exceeded the initial statistical threshold. Details of these clusters are provided in Table 2. The clusters were associated primarily with the orbito-frontal and medial frontal cortex and the anterior cingulate.

#### TS group > CS group contrast

The analysis revealed no clusters that were statistically significant and survived correction for multiple comparisons.

### Relationship between cingulate grey matter volume and clinical measures

To examine the association between the cingulate GM values and clinical measures of tic severity (measured using the YGTSS) and premonitory urges (measured using the PUTS) we identified the spatial location of voxels within the cingulate ROI that were significantly correlated with YGTSS motor tic scores and/or premonitory urge (PUTS) scores. In each case the initial (absolute) correlation threshold was set at r = 0.3 and the statistical threshold was then corrected for multiple comparisons using the false-discovery rate [FDR] with alpha set at p < 0.05 (Benjamini & Hochberg, 1995).

This analysis revealed three clusters of voxels that were significantly associated with one or other of the clinical measures. A left hemisphere posterior cingulate (PCC) cluster and a left hemisphere anterior mid cingulate cluster were identified that were positively associated with YGTSS motor tic severity scores. A left hemisphere anterior mid cingulate (aMCC) cluster was identified that was positively associated with PUTS scores. The spatial distribution of these two clusters is illustrated in Table 3 and illustrated in Figure 2.

**Table 3:** Regions in which GM values within the cingulate ‘seed’ ROI were significantly associated with motor tic severity or premonitory urge scores in individuals with TS (r > 0.3, p < 0.05 FDR-corrected, K (cluster-size >= 20).

| Cluster | Size | r-value | Region | Hemisphere | Peak MNI coordinates |  |  |
| --- | --- | --- | --- | --- | --- | --- | --- |
|  |  |  |  |  | X | Y | Z |
| 1 | 33 | 0.44 | Posterior CC | L | -11 | -9 | 43 |
| 2 | 27 | 0.44 | Anterior Mid CC | L | -11 | 27 | 27 |
|  |  |  |  |  | X | Y | Z |
| 1 | 30 | 0.44 | Posterior Mid CC | L | -5 | 7 | 27 |

**Figure 2:**
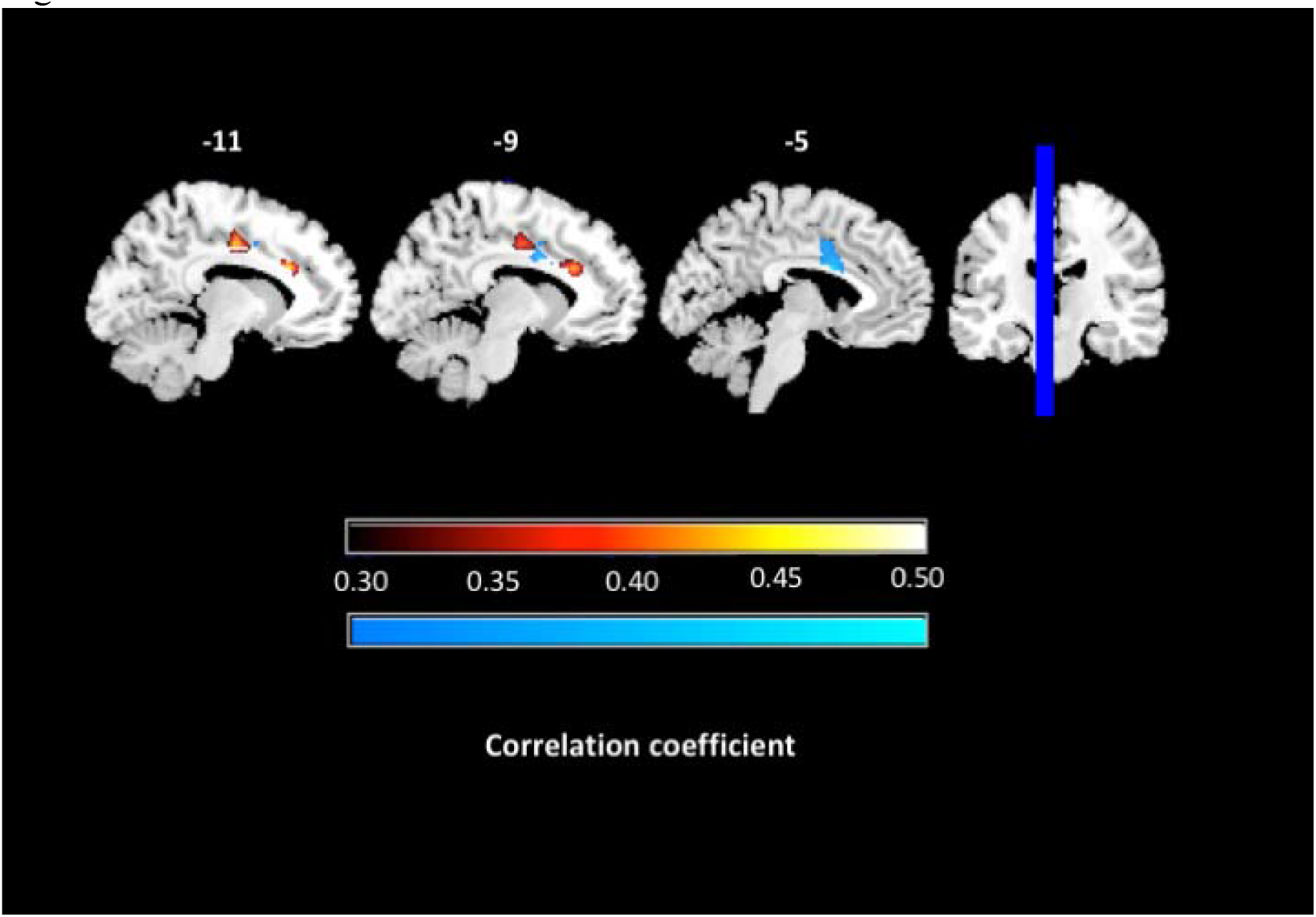
Illustrates clusters of cingulate voxels that were significantly associated with either YGTSS motor tic severity (warm colours) or premonitory urge (PUTS) score (cool colours). These regions consisted of: a posterior cingulate and separate anterior mid cingulate cluster that was positively associated with motor tic severity, and a separate anterior mid cingulate cluster that was significantly positively correlated with premonitory urge (PUTS) scores.

**Figure 3:**
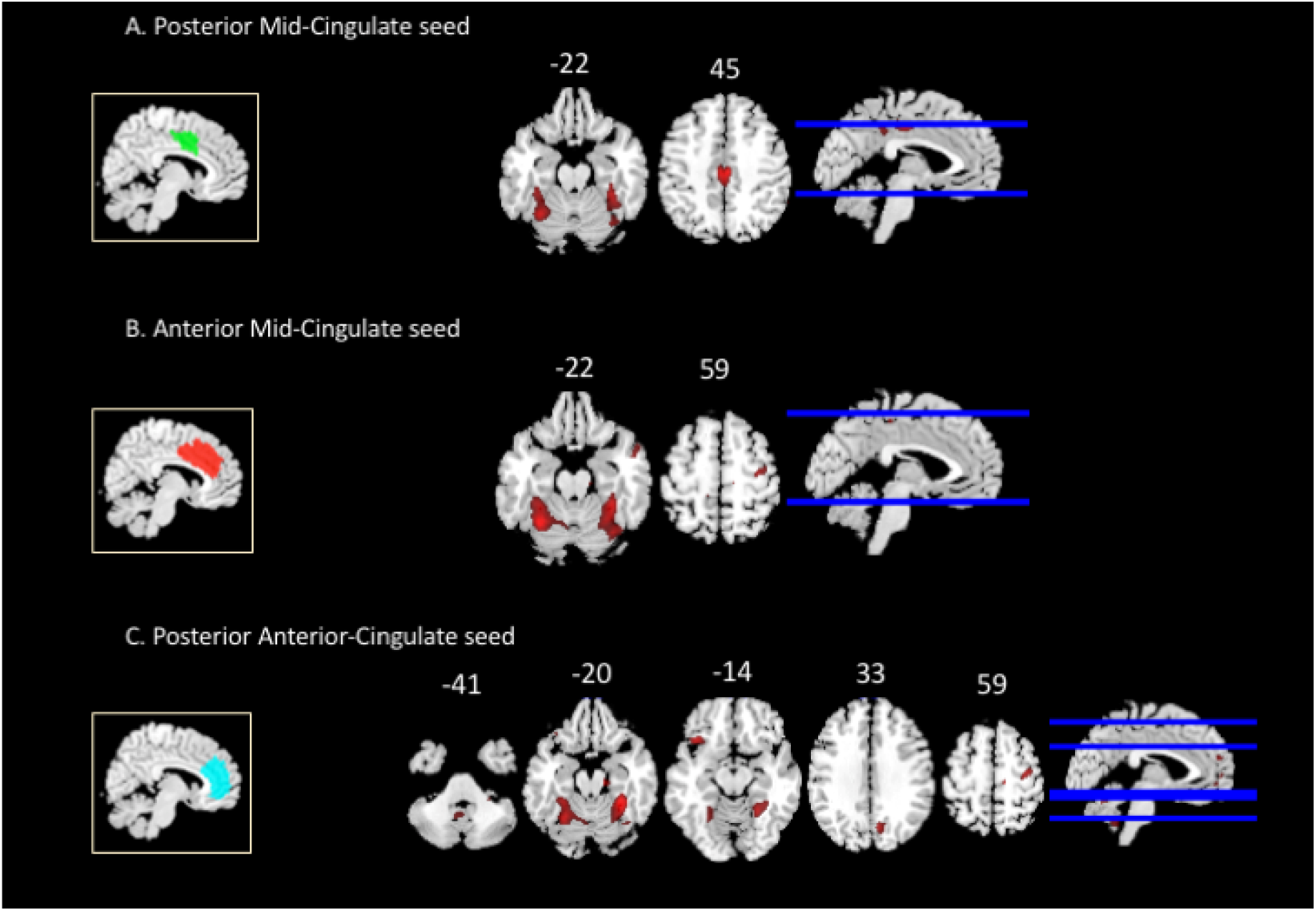
Illustrates the results of the structural covariance analysis for each of the three bilateral cingulate ROIs (shown in the yellow boxes). The upper panel shows clusters of voxels with *increased* structural covariance with the mean GM value of the pMCC ROI region (green) in the TS group relative to the healthy control group. The middle panel shows clusters of voxels with *increased* structural covariance with the mean GM value of the aMCC ROI region (red) in the TS group. The lower panel shows clusters of voxels with *increased* structural covariance with the mean GM value of the pACC ROI region (cyan) in the TS group.

### Structural covariance network analyses

The findings outlined above, together with previous functional brain imaging (e.g., Balsters et al., 2016) and electrical brain stimulation studies (e.g., Caruana et al., 2018) indicate that the cingulate cortex can be partitioned into functionally distinct regions with distinct behavioural characteristics. For this reason, and in order to investigate alterations in structural covariance networks associated with TS, we constructed three functionally-defined bilateral cingulate ROIs based upon the published control data presented in Balsters et al., 2016 (connectivity based probability maps of the cingulate cortex for typically developing young adults are available for download at http://www.ncm.hest.ethz.ch/downloads/data.html). Specifically, based upon our own findings, functional MRI studies investigating the neural antecedents of tics in TS (e.g., Bolhalter et al., 2006; Neuner et al., 2014), MRI-based functional connectivity studies of the cingulate cortex (e.g., Balsters et al., 2016), and electrical stimulation studies of the cingulate cortex (e.g., Caruana et al., 2018), we constructed the following: a bilateral posterior mid-cingulate (pMCC) ROI; a bilateral anterior mid-cingulate (aMCC) ROI, and a posterior (pregunual) anterior cingulate (pACC) ROI as separate structural covariance seed regions.

We then investigated whether there were statistically significant between-group differences in the structural covariance networks (SCNs) linked to each of these seed regions. As noted above (Methods), we did this by comparing the covariance maps for each group (in which the value at each voxel reflected the cross-subject Pearson correlation coefficient between the mean GM value for the seed region and the GM value at that particular voxel) and computing a Z statistic at each voxel which represented the difference between the correlation coefficients. An initial statistical (Z) threshold of 2.5 was adopted and corrected for multiple comparisons using FDR (Benjamini & Hochberg, 1995) together a cluster threshold of K_E_ ≥ 50 voxels). The results of these analyses are outlined below.

#### Posterior mid-cingulate (pMCC)

Analysis of the SCNs associated with the bilateral pMCC revealed four clusters of 50+ voxels in which the structural covariance with the pMCC seed region was significantly *increased* in the TS group relative to the control group. Note that in this case, these clusters exhibited significantly larger correlations with the seed region than was the case for the typically developing controls (i.e., Z difference scores). Details of these clusters are presented in Table 4 (upper panel) and illustrated in Figure 4. They include: the bilateral cerebellum (Lobule VI); the bilateral posterior cingulate cortex and the right inferior temporal cortex. There were no clusters in which the structural covariance with the pMCC seed region was significantly decreased in the TS group.

**Table 4:** Regions in which structural covariance values with the cingulate ‘seed’ ROI were significantly increased in individuals with TS relative to those for typically developing controls (Z > 2.5, p < 0.05 FDR-corrected, K (cluster-size >= 50).

| **TS > CS: posterior mid-cingulate seed** |  |  |  |  |  |  | MNI coordinates |  |  |
| --- | --- | --- | --- | --- | --- | --- | --- | --- | --- |
| Cluster | Size | z-value | CS<br>r(Z) | TS<br>r(Z) | Region | Hemisphere | X | Y | Z |
| 1 | 548 | 3.65 | -0.33 | 0.48 | Cerebellum<br>(Lobule VI) | R | 37 | -67 | -19 |
| 2 | 382 | 3.66 | -0.29 | 0.59 | Cerebellum<br>(Lobule VI) | L | -33 | -61 | -23 |
| 3 | 345 | 4.31 | 0.14 | 0.82 | Post. cingulate<br>cortex | L/R | -5 | -19 | 41 |
| 4 | 64 | 3.30 | 0.02 | 0.49 | Inf. Temporal<br>cortex | R | 21 | -5 | -47 |

| **TS > CS: anterior mid-cingulate seed** |  |  |  |  |  |  | MNI coordinates |  |  |
| --- | --- | --- | --- | --- | --- | --- | --- | --- | --- |
| Cluster | Size | z-value | CS<br>r(Z) | TS<br>r(Z) | Region | Hemisphere | X | Y | Z |
| 1 | 844 | 3.86 | -0.27 | 0.63 | Cerebellum<br>(Lobule VI) | R | 33 | -45 | -21 |
| 2 | 618 | 4.26 | -0.32 | 0.67 | Cerebellum<br>(Lobule VI) | L | -31 | -61 | -23 |
| 3 | 112 | 3.52 | -0.27 | 0.58 | Pre-central<br>Gyrus | R | 47 | -15 | 67 |
| 4 | 54 | 3.47 | -0.19 | 0.41 | Mid. Temporal<br>Gyrus | R | 55 | 11 | -27 |

| **TS > CS: posterior anterior-cingulate seed** |  |  |  |  |  |  | MNI coordinates |  |  |
| --- | --- | --- | --- | --- | --- | --- | --- | --- | --- |
| Cluster | Size | z-value | CS<br>r(Z) | TS<br>r(Z) | Region | Hemisphere | X | Y | Z |
| 1 | 1238 | 4.18 | -0.28 | 0.68 | Cerebellum<br>(Lobule VI) | R | 33 | -45 | -21 |
| 2 | 775 | 3.60 | -0.27 | 0.59 | Cerebellum<br>(Lobule VI) | L | -29 | -61 | -21 |
| 3 | 219 | 3.46 | 0.11 | 0.77 | Mid Frontal<br>Gyrus | R | 39 | -11 | 59 |
| 4 | 147 | 3.38 | -0.16 | 0.63 | Precuneus | R | 11 | -69 | 33 |
| 5 | 138 | 3.69 | -0.32 | 0.47 | Cerebellum<br>(Lobule V) | R | 3 | -65 | -1 |
| 6 | 132 | 3.55 | 0.44 | 0.82 | Sup. Medial<br>Gyrus | L | -13 | 55 | 5 |
| 7 | 112 | 3.65 | -0.12 | 0.69 | Temporal Pole | L | -39 | 21 | -15 |
| 8 | 105 | 2.94 | -0.38 | 0.39 | Cerebellum<br>(Lobule IX) | L | -7 | -59 | -41 |
| 9 | 103 | 3.12 | -0.12 | 0.46 | Cerebellum<br>(Lobule VIIIb) | R | 25 | -39 | -41 |

#### Anterior mid-cingulate (aMCC)

Analysis of the SCNs associated with the bilateral aMCC revealed four clusters of 50+ voxels in which the structural covariance with the aMCC seed region was significantly *increased* in the TS group relative to the control group. Details of these clusters are presented in Table 4 (middle panel) and illustrated in Figure 4. Inspection of Table 4 shows that the clusters of increased covariance with the aMCC seed are highly similar to those identified for the pMCC. They include: the bilateral cerebellum (Lobule VI); the right motor and right temporal cortex. There were no clusters in which the structural covariance with the aMCC seed region was significantly decreased in the TS group.

#### Posterior anterior-cingulate (pACC)

Analysis of the SCNs associated with the bilateral pACC revealed nine clusters of 50+ voxels in which the structural covariance with the pACC seed region was significantly *increased* in the TS group relative to the control group. Details of these clusters are presented in Table 4 (lower panel) and illustrated in Figure 4. Inspection of Table 4 again shows that the clusters of increased covariance associated with the pACC seed are highly similar to those identified for both the pMCC and aMCC. They include: the bilateral cerebellum (Lobules V, VI, VIII, and IX), bilateral frontal lobe, right precuneus, and the left temporal pole. There were no clusters in which the structural covariance with the pACC seed region was significantly decreased in the TS group.

## Discussion

In this study we utilised voxel-based morphometry techniques together with ‘seed-to-voxel’ structural covariance network (SCN) mapping to investigate the putative role played by the mid and anterior cingulate cortex in the generation of motor tics and the experience of PU in a relatively large group of young people TS. Our findings demonstrate the following. First, that global measures of GM volume such as total number of GM voxels or average GM volume are significantly reduced in individuals with a diagnosis of TS, independently of chronological age, sex, or IQ. Second, that whole-brain VBM analysis demonstrate that TS is associated primarily with significant reduction in GM volume within the orbito-frontal cortex and the the anterior and mid cingulate cortex. Third, that clinical measures of motor tic severity was positively associated with two separate clusters of voxels within the left hemisphere posterior cingulate cortex (PCC) and anterior mid cingulate cortex (aMCC). Whereas clinical measures of premonitory urge (PUTS) were associated with a left hemisphere anterior mid cingulate (aMCC) cluster. Fourth, when we examined the structural covariance networks (SCNs) associated with three bilateral regions of the cingulate cortex (pMCC, aMCC and pACC), we found that these networks differed in individuals with TS compared to the group of typically developing individuals. In each case we found that there was increased structural covariance between the cingulate cortex ROI and a set of brain areas that included in each case: the bilateral motor cerebellum (Lobule VI); the posterior cingulate cortex; and inferior frontal cortex. There were no regions identified in which structural covariance with the cingulate was reduced for the TS group.

### Reduction in GM volume

The finding that there are widespread decreases in GM volume in individuals with TS has reported on many previous studies (e.g., Draganski, Martino, Cavanna, Hutton, Orth, Robertson, et al. 2010; Draper, Jackson, Morgan, Jackson, 2016; Fahim C, Yoon U, Sandor P, et al., 2009; Greene et al., 2017; Müller-Vahl, Kaufmann, Grosskreutz, et al., 2009; Peterson, Pine, Cohen, et al., 2001; Sowell, Kan, Yoshii, et al., 2008; Worbe, Gerardin, Hartmann, et al., 2010), and the results of the current study confirm this finding. Two aspects of our current findings are worthy of note. First, it has been suggested previously that alterations in brain structure during adolescence in TS - such as reduced GM volume or decreased WM connectivity - might reflect neuroplastic adaptations that result in a reduction in clinical symptoms such as tic severity (e.g., Jackson, Parkinson, Jung, Ryan, Morgan, Hollis et al., 2011; Plessen, Bansal, Peterson, 2009; Plessen et al., 2004). In the current study we find that the rate at which GM volume decreases with chronological age, as indexed by the total number of GM voxels identified within each individual’s GM map, is greater for the TS group compared to the healthy controls, and that this effect is independent of chronological age, sex or IQ. This finding is consistent with that previously reported by Draganski et al., (2010), who demonstrated that GM volume within the ACC decreases faster with age in their TS group compared to healthy controls. One might speculate that the increased rate at which GM volume decreases in TS over adolescence might well reflect increased synaptic pruning or a related developmental process. However, this question is best answered by a longitudinal investigation of how brain structure changes over adolescence, and how such changes are related to clinical symptoms in TS.

Second, our VBM analysis demonstrated that for the TS group there were decreases in GM volume primarily within: the orbito-frontal, medial frontal, and motor cortices and within the cingulate cortex (all p < 0.001^uncorrected^). However, none of these clusters survived statistical correction for multiple comparison, and must therefore be interpreted with this in mind. These brain regions have previously been particularly associated with the pathophysiology of tics in TS (see Cavanna et al., 2017, O’ Neill et al. 2019, Pedroarena-Leal & Ruge, 2015, and Plessen et al., 2009 for reviews) and our results confirm previous MRI studies that report structural alteration in TS within: the orbito-frontal cortex (e.g., Draganski et al., 2010; Greene et al., 2017); and the MCC and pACC regions of the cingulate cortex (e.g., Draganski et al., 2010; Müller-Vahl et al., 2009; Worbe et al., 2010). It is noteworthy that electrical stimulation of the pMCC, aMCC and pACC regions of the cingulate cortex is sufficient to initiate goal-directed movements in humans (Caruana et al., 2018).

### Anatomical separation of clinical measures of tic severity and PU within the cingulate cortex

The cingulate cortex has long been divided into different anatomical regions. Most schemes separate the cingulate cortex into anterior (ACC), mid (MCC) and posterior (PCC) regions, but the ACC and MCC have been further divided into posterior (pre-genual) and anterior (sub-genual) regions for the ACC, and posterior and anterior areas for the MCC (Palomero-Gallagher, Eickhoff, Hoffstaedter, Schleicher, Mohlberg, Vogt, et al., 2015; Palomero-Gallagher, Mohlberg, Zilles, Vogt, 2008; Palomero-Gallagher, Vogt, Schleicher, Mayberg, Zilles, 2009). However, the functional roles of the cingulate, including any functional specialisation of anatomically defined sub-regions, has been less clear.

In the current study we conducted a correlation analysis which demonstrated that there were clusters of voxels in the posterior cingulate and anterior mid-cingulate cortex that were positively associated with motor tic severity scores, and a separate cluster within the anterior mid-cingulate cortex that was positively correlated with premonitory urge scores. Our finding that clusters of GM volume of voxels within the cingulate cortex are associated with clinical measures of motor tic severity and premonitory urge scores in TS is consistent with the proposed involvement of the cingulate cortex in the generation of tics in response to premonitory urges (e.g., Devinsky, 1983, 1995; Jackson, Parkinson, Kim, et al., 2011) and with the finding that the mid-cingulate cortex is activated ahead of tic execution in individuals with TS (Bolhalter et al, 2006, Neuner et al., 2014). Recent electrical stimulation studies of human cingulate cortex have demonstrated functional differences between the PCC, MCC and ACC. In one study, stimulation of the pre-genual region of the ACC produced emotional, interoceptive and autonomic responses, whereas stimulation of the MCC, particularly the anterior region, produced goal-directed movements. By contrast, the PCC and was generally unresponsive in this study when stimulated (Caruana, et al., 2018). However, in another study of electrical stimulation of the medial wall in humans, it was found that stimulation of the caudal cingulate zone (corresponding to the posterior MCC) produced primarily somatosensory responses (primarily paraesthesias, dysesthesias, or pain) but not overt movement (Trevisi et al., 2018). These findings are broadly consistent with our finding that: reduction of GM volume in the MCC region is associated with increased motor tic severity and premonitory urge scores; increases in GM volume in the PCC and aMCC were associated with increased motor tic severity; and increases in GM volume in the aMCC region were associated with increased premonitory sensory phenomena.

### Between-group differences in structural covariance networks (SCNs) associated with the functionally defined the MCC and ACC

As noted above, previous studies have reported that structural covariance networks (SCNs) closely mirror the intrinsic functional connectivity networks identified using resting-state fMRI (e.g., (Kelly et al., 2012; Seeley et al., 2009), and it has been proposed that SCNs likely reflect brain areas that are functionally co-active and subject to common patterns of maturational change and gene co-expression (Romero-Garcia et al., 2018; Zielinski et al., 2010). Such SCNs are selectively vulnerable to specific brain health conditions (Seeley et al., 2009) and co-degenerate in specific neurodegenerative conditions (Cauda et al., 2018; Seeley et al., 2009).

In their urge-for-action model, Jackson et al. (2011) proposed that signals associated with the bodily experienced as PU in TS are relayed from the insula to the cingulate motor areas that, together with the ventral striatum, may participate in the selection of a particular action based upon an analysis of the likely “value” of that action given the organism’s previous history of action outcomes. This proposal is consistent with reports that the cingulate motor areas (MCC) is activated immediately prior to the execution of tics in TS (e.g., Bohlhalter et al., 2006; Neuner et al., 2014) and that electrical stimulation of the MCC region of the cingulate cortex is sufficient to induce the execution of goal-directed movements (e.g., Caruana, et al., 2018).

In the current study we compared the structural covariance networks (SCNs) associated with three functionally-defined bilateral regions of the cingulate cortex (pMCC, aMCC and pACC) in our TS group to those of a group of typically developing controls (having first controlled for the effects of chronological age, TIV, IQ and sex). We found that each of these SCNs differed in individuals with TS compared to the group of typically developing individuals in quite similar ways. Specifically, we found that in each case there was *increased* structural covariance between the cingulate cortex ROI and a set of brain areas that included: the bilateral motor cerebellum (Lobule VI); the posterior cingulate cortex; the inferior temporal cortex. There were no regions identified in which structural covariance was reduced for the TS group.

Given the putative role outlined above for the cingulate cortex in generating movements in response to premonitory urge signals most likely arising from the right insula cortex (Jackson, Parkinson, Kim, et al., 2011), it is not a surprise that the TS group exhibit significantly increased structural covariance with motor regions of the cerebellum bilaterally. It is noteworthy that recent physiological recording studies have indicated that the cerebellum may play a key role in gating the expression of tic-like movements in a non-human primate model of motor tics in TS (McCairn et al., 2013), and that structural brain imaging studies of TS have reported significantly reduced GM volume relative to healthy controls in the cerebellum (e.g., Tobe, Bansal, Xu, Hao, Liu, Sanchez, Peterson, 2010) and cingulate cortex (e.g., Müller-Vahl et al., 2009; for a recent review see O’Neill et al., 2019).

To the extent that SCNs have been demonstrated to closely mirror the intrinsic functional connectivity networks identified using resting-state fMRI (e.g., (Kelly et al., 2012; Seeley et al., 2009), and may reflect brain areas that are functionally co-active and subject to common patterns of maturational change and gene co-expression (Romero-Garcia et al., 2018; Zielinski et al., 2010), it is tempting to speculate that the increased structural covariance between cingulate and motor cerebellum represents long-term increased functional connectivity of this urge-for-action network in individuals with TS. However, we recognise that this proposal can best be investigated by looking directly at functional connectivity between these areas using functional brain imaging techniques.

In the current study we found increased structural covariance between the cingulate cortex and the inferior frontal cortex in TS. It is widely believed that the prefrontal cortex, and in particular the right inferior frontal cortex, plays a key role in the inhibitory control of other brain regions (Aron, 2007), including brain areas associated with the urge-for-action (Berman, Horovitz, Morel, Hallett, 2012). Consistent with this view, Berman et al. (2012) demonstrated that increased brain activity within the right inferior frontal cortex, bilateral insula and mid-cingulate cortex were associated with a build-up of the urge-to-blink, and they concluded that the right inferior frontal cortex may function to maintain volitional control over motor output in spite of an increasing sense of urge for action (Berman et al., 2012).

It has been suggested that individuals with TS may gain control over their tics through the development of compensatory self-regulation mechanisms, implemented through changes in neural pathways linking inhibitory control regions of prefrontal cortex with primary and secondary motor areas (e.g., Peterson et al., 2001; Serrien, et al., 2005; Plessen et al., 2009; Jackson, Parkinson, Jung, et al., 2011; Jackson, Draper, Dyke, Pépés, Jackson, 2015). Consistent with this proposal, studies of cognitive control of motor outputs demonstrated that individuals with ‘uncomplicated’ TS (i.e., those without co-morbid disorders such as ADHD) exhibit paradoxically *enhanced* volitional control over their motor behaviour (Mueller, Jackson, Dhalla, Datsopoulos, Hollis, 2006; Jackson, Mueller, Hambleton, Hollis, 2007; Jackson, Parkinson, Jung, et al., 2011) and reduced corticospinal excitability ahead of volitional movements (Heise et al., 2010; Jackson, Parkinson, Manfredi, Millon, Hollis, Jackson, 2013; Draper, Stephenson, Jackson, Pépés, Morgan, Morris, et al., 2014). These findings are consistent with the proposal that the frequent need to actively suppress tics leads to a generalised enhancement in the efficacy of volitional control mechanisms in TS. Our current finding of increased structural covariance between inferior frontal and anterior cingulate cortex may represent increased long-term functional connectivity associated with volitional suppression of the cingulate urge-for-action network in individuals with TS.

#### Limitations of this study

Throughout this paper we have assumed that structural covariance networks (SCNs) closely mirror the intrinsic connectivity networks (ICNs) measured and robustly demonstrated using resting-state fMRI. This assumption is based upon a number of studies, referred to above, that have directly compared seed-based SCNs and ICNs. However, given that our study investigates SCNs in adolescents and young adults with TS – a period during which the brain is known to undergo considerable maturation – it is possible that during this period there could be differences in the maturation of structural and functional brain networks, and that these may be exacerbated by the presence of TS. For this reason, as we have not directly measured functional brain connectivity in this study, we advocate caution in drawing strong conclusions about functional connectivity based upon our findings of structural covariance differences in TS.

While the majority of our sample of adolescents and young adults with TS were unmedicated, we recognise that we are unable to say very little about the effects of medication on our findings. Thus, while we have controlled for variables such as age, sex, IQ and intracranial volume in our analyses, we were unable to investigate the effects of medication or co-morbidities due to the heterogeneous nature of the sample, and we have interpreted our results with this in mind.

#### Conclusion

In summary, in the current study we used voxel-based morphometry together with ‘seed-to-voxel’ structural covariance mapping to investigate the role played by the cingulate cortex in the generation of motor tics and the experience of PU in a group of young people TS. We demonstrated that global measures of GM volume such as total number of GM voxels or average GM volume are significantly reduced in individuals with a diagnosis of TS, independently of chronological age, sex, or IQ, and that TS is associated with reduction in GM volume particularly within the cerebellum and the cingulate cortex. We also demonstrated that motor tic severity and PU scores were associated with anatomically separate regions of the cingulate cortex. Specifically, an posterior cingulate cluster and anterior mid cingulate cluster that was positively associated with motor tic severity; and an anterior mid cingulate cluster that was positively associated with premonitory urge scores. Finally, we examined the structural covariance networks (SCNs) associated with three bilateral regions of the cingulate cortex (pMCC, aMCC and pACC) and found that these networks differed in individuals with TS compared to the group of typically developing individuals. Specifically, we found increased structural covariance between the cingulate cortex ROI and a set of brain areas that included in each case the bilateral motor cerebellum (Lobule VI), the posterior cingulate cortex, and in the case of the ACC only, the inferior frontal cortex. This suggests that analysis of SCNs, while currently under-utilised in the study of brain networks in neurodevelopmental conditions, may be a particularly useful method for investigating alterations in brain network development in children and adolescents, in particular for those for whom the use of conventional fMRI approaches is especially challenging.

## Acknowledgements

This work was supported by the Medical Research Council (grant number G0901321), the James Tudor Foundation, Tourettes Action (UK), and by the NIHR Nottingham Biomedical Research Centre. The views expressed are those of the authors and not necessarily those of the NHS, the NIHR or the Department of Health. The authors would like to thank Jane Fowlie and Tourettes Action (UK) for assisting with participant recruitment.

## References

Alexander-Bloch, A., Giedd, J. N., & Bullmore, E. T. (2013). Imaging structural co-variance between human brain regions. Nature Reviews Neuroscience, 14(5), 322–336. doi: 10.1038/nrn3465

Aron, A. R. (2007). The neural basis of inhibition in cognitive control. Neuroscientist, 13(3), 214–228. doi: 10.1177/10738584072992

Aron, A.R., Robbins, T.W., and Poldrack, R.A. (2004). Inhibition and the right inferior frontal cortex. Trends Cogn. Sci. 8, 170–177.

Ashburner, J. (2007). A fast diffeomorphic image registration algorithm. Neuroimage, 38(1), 95–113. doi: 10.1016/j.neuroimage.2007.07.007

Balsters JH, Mantini D, Apps MAJ, Eickhoff SB, Wenderoth N. (2016). Connectivity-based parcellation increases network detection sensitivity in resting state fMRI: An investigation into the cingulate cortex in autism. NeuroImage: Clinical 11, 494–507.

Benjamini, Y., & Hochberg, Y. (1995). Controlling the False Discovery Rate - a Practical and Powerful Approach to Multiple Testing. Journal of the Royal Statistical Society Series B-Statistical Methodology, 57(1), 289–300. doi: 10.1111/j.2517-6161.1995.tb02031

Berman, B. D., Horovitz, S. G., Morel, B., & Hallett, M. (2012). Neural correlates of blink suppression and the buildup of a natural bodily urge. Neuroimage, 59(2), 1441–1450. doi: 10.1016/j.neuroimage.2011.08.05

Berument, S. K., Rutter, M., Lord, C., Pickles, A., & Bailey, A. (1999). Autism screening questionnaire: diagnostic validity. British Journal of Psychiatry, 175, 444–451. doi: 10.1192/bjp.175.5.444

Bohlhalter, S., Goldfine, A., Matteson, S., Garraux, G., Hanakawa, T., Kansaku, K., … Hallett, M. (2006). Neural correlates of tic generation in Tourette syndrome: an event-related functional MRI study. Brain, 129(8), 2029–2037. doi: 10.1093/brain/awl050

Bronfeld, M., Israelashvili, M., & Bar-Gad, I. (2013). Pharmacological animal models of Tourette syndrome. Neuroscience and Biobehavioral Reviews, 37(6), 1101–1119. doi: 10.1016/j.neubiorev.2012.09.010

Caruana F, Gerbella M, Avanzini P, Gozzo F, Pelliccia V, Mai R, Abdollahi RO, Cardinale F, Sartori I, Lo Russo G, Rizzolatti G, (2018). Motor and emotional behaviours elicited by electrical stimulation of the human cingulate cortex. Brain 141, 3035–3051.

Cauda, F., Nani, A., Manuello, J., Premi, E., Palermo, S., Tatu, K., … Costa, T. (2018). Brain structural alterations are distributed following functional, anatomic and genetic connectivity. Brain, 141(11), 3211–3232. doi: 10.1093/brain/awy252

Cavanna, A. E., Black, K. J., Hallett, M., & Voon, V. (2017). Neurobiology of the premonitory urge in Tourette’s syndrome: pathophysiology and treatment implications. J Neuropsych Clin N, 29(2), 95–104. https://doi.org/10.1176/appi.neuropsych.16070141

Cohen, S. C., Leckman, J. F., & Bloch, M. H. (2013). Clinical assessment of Tourette syndrome and tic disorders. Neuroscience Biobehavioural Reviews, 37(6), 997–1007. doi: 10.1016/j.neubiorev.2012.11.013

Cohrs, S., Rasch, T., Altmeyer, S., Kinkelbur, J., Kostanecka, T., Rothenberger, A., … Hajak, G. (2001). Decreased sleep quality and increased sleep related movements in patients with Tourette's syndrome. Journal of Neurology, Neurosurgery and Psychiatry, 70(2), 192–197. doi: 10.1136/jnnp.70.2.192.

Conceicao, V. A., Dias, A., Farinha, A. C., & Maia, T. V. (2017). Premonitory urges and tics in Tourette syndrome: computational mechanisms and neural correlates. Current Opinions in Neurobiology, 46, 187–199. doi: 10.1016/j.conb.2017.08.009

Conners, C. K. (2008). Conners third edition (Conners 3). Los Angeles, CA: Western Psychological Services.

Devinsky O (1983). Neuroanatomy of Gilles de la Tourette’s syndrome: possible midbrain involvement. Arch Neurol 40: 508–514.

Devinsky O, Morrell MJ, Vogt BA (1995). Contributions of anterior cingulate to behavior. Brain 118: 279–306.

Draganski, B., Martino, D., Cavanna, A. E., Hutton, C., Orth, M., Robertson, M. M., … Frackowiak, R. S. (2010). Multispectral brain morphometry in Tourette syndrome persisting into adulthood. Brain, 133, 3661–3675. doi: 10.1093/brain/awq300

Draper, A., Jackson, G. M., Morgan, P. S., & Jackson, S. R. (2016). Premonitory urges are associated with decreased grey matter thickness within the insula and sensorimotor cortex in young people with Tourette syndrome. Journal of Neuropsychology, 10(1), 143–153. doi: 10.1111/jnp.12089

Draper A, Stephenson MC, Jackson GM, Pépés S, Morgan PS, Morris PG, Jackson SR (2014). Increased GABA Contributes to Enhanced Control over Motor Excitability in Tourette Syndrome, Current Biology 24: 2343–2347.

Eickhoff, S. B., Stephan, K. E., Mohlberg, H., Grefkes, C., Fink, G. R., Amunts, K., & Zilles, K. (2005). A new SPM toolbox for combining probabilistic cytoarchitectonic maps and functional imaging data. Neuroimage, 25(4), 1325–1335. doi: 10.1016/j.neuroimage.2004.12.034

Fahim C, Yoon U, Sandor P et al. (2009). Thinning of the motor–cingulate–insular cortices in siblings concordant for Tourette syndrome. Brain Topogr 22: 176–184.

Fried, I., Katz, A., McCarthy, G., Sass, K. J., Williamson, P., & Spencer, S. S. (1991). Functional organization of human supplementary motor cortex studied by electrical stimulation. The Journal of Neuroscience, 11(11), 3656e3666.

Ganos, C., Garrido, A., Navalpotro-Gómez, I., Ricciardi, L., Martino, D., Edwards, M. J., … Bhatia, K. P. (2015). Premonitory Urge to Tic in Tourette’s Is Associated With Interoceptive Awareness. Movement Disorders, 30(9), 1198–1202. doi: 10.1002/mds.26228

Ganos, C., Kahl, U., Schunke, O., Kuhn, S., Haggard, P., Gerloff, C., … Munchau, A. (2012). Are premonitory urges a prerequisite of tic inhibition in Gilles de la Tourette syndrome? Journal of Neurology Neurosurgery and Psychiatry, 83(10), 975–978. doi: 10.1136/jnnp-2012-303033

Gratton, C., Laumann, T. O., Nielsen, A. N., Greene, D. J., Gordon, E. M., Gilmore, A. W., … Petersen, S. E. (2018). Functional Brain Networks Are Dominated by Stable Group and Individual Factors, Not Cognitive or Daily Variation. Neuron, 98(2), 439–452. doi: 10.1016/j.neuron.2018.03.035

Greene, D. J., Williams Iii, A. C., Koller, J. M., Schlaggar, B. L., Black, K. J., & The Tourette Association of America Neuroimaging, C. (2017). Brain structure in pediatric Tourette syndrome. Molecular psychiatry, 22(7), 972–980. doi: 10.1038/mp.2016.194

Heise, K.F. et al. (2010) Altered modulation of intracortical excitability during movement preparation in Gilles de la Tourette syndrome. Brain 133, 580–590

Jackson GM, Draper A, Dyke K, Pépés SE, Jackson SR. (2015). Inhibition, disinhibition and the control of action in Tourette syndrome. Trends in Cognitive Sciences 19: 655–665.

Jackson, G.M., Mueller, S.C., Hambleton, K., and Hollis, C.P. (2007). Enhanced cognitive control in Tourette Syndrome during task uncertainty. Exp. Brain Res. 182, 357–364.

Jackson, S. R., Parkinson, A., Jung, J., Ryan, S. E., Morgan, P. S., Hollis, C., & Jackson, G. M. (2011). Compensatory neural reorganization in Tourette syndrome. Current Biology, 21(7), 580–585. doi: 10.1016/j.cub.2011.02.047

Jackson, S. R., Parkinson, A., Kim, S. Y., Schuermann, M., & Eickhoff, S. B. (2011). On the functional anatomy of the urge-for-action. Cognitive Neuroscience, 2(3-4), 227–243. doi: 10.1080/17588928.2011.604717

Jackson SR, Parkinson A, Manfredi V, Millon G, Hollis CP, Jackson GM. (2013). Motor excitability is reduced prior to voluntary movements in children and adolescents with Tourette syndrome. Journal of Neuropsychology 7: 29–44.

Kelly, C., Toro, R., Di Martino, A., Cox, C. L., Bellec, P., Castellanos, F. X., & Milham, M. P. (2012). A convergent functional architecture of the insula emerges across imaging modalities. Neuroimage, 61(4), 1129–1142. doi: 10.1016/j.neuroimage.2012.03.021

Leckman, J.F., Riddle, M.A., Hardin, M.T., Ort, S.I., Swartz, K.L., Stevenson, J., and Cohen, D.J. (1989). The Yale Global Tic Severity Scale: Initial testing of a clinician-rated scale of tic severity. J. Am. Acad. Child Adolesc. Psychiatry 28, 566–573.

McCairn, K. W., Iriki, A., & Isoda, M. (2013). Global dysrhythmia of cerebro-basal ganglia–cerebellar networks underlies motor tics following striatal disinhibition. The Journal of Neuroscience, 33(2), 697–708. doi: 10.1523/JNEUROSCI.4018-12.2013

Mueller, S.C., Jackson, G.M., Dhalla, R., Datsopoulos, S., and Hollis, C.P. (2006). Enhanced cognitive control in young people with Tourette’s syndrome. Current Biology 16, 570–573.

Müller-Vahl KR, Kaufmann J, Grosskreutz J et al. (2009). Prefrontal and anterior cingulate cortex abnormalities in Tourette syndrome: evidence from voxel-based morphometry and magnetization transfer imaging. BMC Neurosci 10: 47.

Müller-Vahl, K. R., Riemann, L., & Bokemeyer, S. (2014). Tourette patients’ misbelief of a tic rebound is due to overall difficulties in reliable tic rating. Journal of psychosomatic research, 76(6), 472–476. doi: 10.1016/j.jpsychores.2014.03.003

Neuner, I., Werner, C. J., Arrubla, J., Stoecker, T., Ehlen, C., Wegener, H. P., … Shah, N. J. (2014). Imaging the where and when of tic generation and resting state networks in adult Tourette patients. Frontiers in Human Neuroscience, 8. doi: 10.3389/fnhum.2014.00362

O’Neill J, Piacentini JC, Peterson BS. (2019). In Handbook of Clinical Neurology, Vol. 166 (3rd series) Cingulate Cortex B.A. Vogt, Editor. https://doi.org/10.1016/B978-0-444-64196-0.00011-X

Palomero-Gallagher N, Eickhoff SB, Hoffstaedter F, Schleicher A, MohlbergH, Vogt BA, Amunts K, Zilles K (2015) Functional organization of human subgenual cortical areas: relationship between architectonical segregation and connectional heterogeneity. Neuroimage 115:177–190.

Palomero-Gallagher N, Mohlberg H, Zilles K, Vogt B (2008) Cytology and receptor architecture of human anterior cingulate cortex. J Comp Neurol 508:906–926.

Palomero-Gallagher N, Vogt BA, Schleicher A, Mayberg HS, Zilles K. (2009) Receptor architecture of human cingulate cortex: Evaluation of the four-region neurobiological model. Human Brain Mapping 30, 2336–2355.

Pedroarena-Leal, N. & Ruge, D. Cerebellar neurophysiology in Gilles de la Tourette syndrome and its role as a target for therapeutic intervention. J. Neuropsychol 11, 327–346 (2017).

Peterson BS, Pine DS, Cohen P, Brook JS. (2001). Prospective, longitudinalstudy of tic, obsessive-compulsive, and attention-deficit/hyperactivitydisorders in an epidemiological sample. J Am Acad Child AdolescPsychiatry 40, 685–95.

Plessen, K. J., Bansal, R., & Peterson, B. S. (2009). Imaging evidence for anatomical disturbances and neuroplastic compensation in persons with Tourette syndrome. Journal of psychosomatic research, 67(6), 559–573. doi: 10.1016/j.jpsychores.2009.07.005

Plessen KJ, Wentzel-Larsen T, Hugdahl K, Feineigle P, Klein K, Staib LH, Leckman JF, Bansal R, Peterson BS. (2004). Altered interhemispheric connectivity in individuals with Tourette’s disorder. American Journal of Psychiatry 161, 2028–2037.

Romero-Garcia, R., Whitaker, K. J., Vasa, F., Seidlitz, J., Shinn, M., Fonagy, P., … Consortium, N. (2018). Structural covariance networks are coupled to expression of genes enriched in supragranular layers of the human cortex. Neuroimage, 171, 256–267. doi: 10.1016/j.neuroimage.2017.12.060

Scahill, L., Riddle, M. A., McSwigginHardin, M., Ort, S. I., King, R. A., Goodman, W. K., … Leckman, J. F. (1997). Children’s Yale-Brown obsessive compulsive scale: Reliability and validity. Journal of the American Academy of Child & Adolescent Psychiatry, 36(6), 844–852. doi: 10.1097/00004583-199706000-00023

Seeley, W. W., Crawford, R. K., Zhou, J., Miller, B. L., & Greicius, M. D. (2009). Neurodegenerative Diseases Target Large-Scale Human Brain Networks. Neuron, 62(1), 42–52. doi: 10.1016/j.neuron.2009.03.024

Serrien, D. J., Orth, M., Evans, A. H., Lees, A. J., & Brown, P. (2005). Motor inhibition in patients with Gilles de la Tourette syndrome: Functional activation patterns as revealed by EEG coherence. Brain, 128, 116–125.

Sowell, E. R., Kan, E., Yoshii, J., Thompson, P. M., Bansal, R., Xu, D., … Peterson, B. S. (2008). Thinning of sensorimotor cortices in children with Tourette syndrome. Nature Neuroscience, 11(6), 637.

Steinberg, T., Baruch, S. S., Harush, A., Dar, R., Woods, D., Piacentini, J., & Apter, A. (2010). Tic disorders and the premonitory urge. Journal of Neural Transmission, 117(2), 277–284. doi: 10.1007/s00702-009-0353-3

Tobe, R. H., Bansal, R., Xu, D., Hao, X., Liu, J., Sanchez, J., & Peterson, B. S. (2010). Cerebellar morphology in Tourette syndrome and obsessive-compulsive disorder. Annals of Neurology, 67(4), 479–487. doi: 10.1002/ana.21918

Trevisi G, Eickhoff SB, Chowdhury F, Jha A, Rodionov R, Nowell M, Miserocchi A, McEvoy AW, Nachev P, Diehl B. (2018). Probabilistic electrical stimulation mapping of human medial frontal cortex. Cortex 109, 336–346

Wechsler, D. (2011). Wechsler Abbreviated Scale of Intelligence–Second Edition (WASI-II). San Antonio, TX: Pearson.

Woods, D. W., Piacentini, J., Himle, M. B., & Chang, S. (2005). Premonitory urge for tics scale (PUTS): Initial psychometric results and examination of the premonitory urge phenomenon in youths with tic disorders. Journal of Developmental and Behavioral Pediatrics, 26(6), 397–403. doi: 10.1097/00004703-200512000-00001

Worbe, Y., Gerardin, E., Hartmann, A., Valabregue, R., Chupin, M., Tremblay, L., … Lehericy, S. (2010). Distinct structural changes underpin clinical phenotypes in patients with Gilles de la Tourette syndrome. Brain, 133(2), 649–3660. doi: 10.1093/brain/awq293

Zielinski, B. A., Gennatas, E. D., Zhou, J. A., & Seeley, W. W. (2010). Network-level structural covariance in the developing brain. Proceedings of the National Academy of Sciences of the United States of America, 107(42), 18191–18196. doi: 10.1073/pnas.1003109107

